# Tomato spotted wilt virus facilitates non-vector spider mite species (*Tetranychus urticae* and *Tetranychus evansi*) on whole tomato plants

**DOI:** 10.1101/2024.09.18.613733

**Authors:** Vandana Gupta, Sarah Grosjean, Aloyse Moreau, David Carbonell, Alison B. Duncan

## Abstract

Coinfections where hosts harbour more than one parasite species are common in nature. Facilitation among parasites enabling them to better exploit host resources is widespread with direct consequences on their life-history. Plant viruses can facilitate their vectors to increase their transmission, but equivalent studies in virus-non vectors are lacking. Here we study facilitation of two (non-vector) species of *Tetranychus* spider mites, *Tetranychus urticae* and *T. evansi*, by a plant virus, tomato spotted wilt virus (TSWV), in coinfections on a host tomato plant, *Solanum lycopersicum*. We compared the effect of different strains of TSWV on life-history traits of *Tetranychus* mites on cut leaves and whole plants of different tomato varieties. TSWV facilitated both species of spider mite on two different tomato varieties on whole plants, with more offspring of both species becoming adults. In contrast, on cut leaves, facilitation of *T. urticae* was much more variable depending on the experiment, viral strain and tomato variety. We attribute this to the non-homogeneous spread of virus throughout the host plant. Virus is first transported to the top leaves, and not middle leaves which we had used for the experiment. Indeed, if facilitation is indirect via the host immune system or resource based via the release of free-amino acids then this may be less efficient on cut plant parts. These results highlight that facilitation among parasites may increase parasite fitness and at the same time levels of virulence experienced by the host.

## Introduction

Coinfections can have both positive or negative effects on parasite life-history (Balmer et al., 2009; McQueen and McKenzie, 2006; Zilio and Koella, 2020) and epidemics (Johnson and Buller, 2011; Rodgers and Bolnick, 2023). Competition among parasites in coinfections is most commonly assumed, and has been well investigated both theoretically and empirically (Alizon et al., 2013; Read and Taylor, 2001). For instance, (Budischak et al., 2018) show that resource competition with hookworms can decrease *Plasmodium culex* density more than 2 fold in humans. In contrast, facilitation (positive) among parasites has been comparatively ignored despite being common (Dallas et al., 2019; Telfer et al., 2010; Zélé et al., 2018).

Theoretical studies show that facilitation among parasites can increase within-host growth (Eswarappa et al., 2012) onward transmission to new hosts (Alizon et al., 2013) and coinfection prevalence (Kamiya et al., 2018). These predictions have been corroborated by experimental studies showing facilitation (Zélé et al., 2018). Facilitation among parasites can occur via multiple mechanisms including the production of public goods aiding the growth of other parasites (Ford et al., 2016) or interactions mediated via the host immune system, for example, negative immune crosstalk or immunosuppression (Dagostin et al., 2023; Graham, 2008; Lello et al., 2018). This latter occurs in sheep whereby suppression of the immune response by the nematode *Haemonchus contortus* facilitates the survival of the nematode *Trichostrongylus colubriformis* (Lello et al., 2018). Also, in spider mites, coinfection with *Tetranychus evansi* which suppresses immune responses can increase the fecundity of *Tetranychus urticae* (Alba et al., 2015). Thus facilitation can impact parasite life-history across different host and parasite systems in nature.

Plants are host to diverse parasites -fungi, bacteria, viruses and insects - and often harbour coinfections. Plant viruses notoriously infect a wide range of plant hosts, and the majority are transmitted by arthropod vectors (Nault, 1997). Virus-vector-plant relationships have been examined in a number of systems and it has been shown that plant viruses can alter the behaviour and phenotype of both their vectors and plant hosts (Fang et al., 2013; Mauck et al., 2010a; Pan et al., 2021) in seemingly adaptive ways to increase transmission (Blanc and Michalakis, 2016). Virus infection can change plant physiology, by modifying the release of volatiles, visual cues and/or nutritional profile (Eigenbrode et al., 2002; Guo et al., 2019; Mauck et al., 2014; Pan et al., 2021). In turn, vectors are often more attracted to virus infected plants (Eigenbrode et al., 2002; Ingwell et al., 2012), and can have increased fitness following feeding on virus infected plants (Chen et al., 2013; Maluta et al., 2014; Maris et al., 2004) but see (Mauck et al., 2010a, 2010b, 2014). Viruses manipulating plant phenotypes can also impact other non-vector parasites sharing the host plant (Ángeles-López et al., 2018; Belliure et al., 2010; Mauck et al., 2010b, 2015). This includes positive effects of viruses on vector species being hijacked by other non-vector parasites sharing the plant (Ángeles-López et al., 2018; Belliure et al., 2010; Nachappa et al., 2013).

Plant hosts have evolved a suite of counterattack mechanisms involving physiological and immune adaptations against parasites (Glazebrook, 2005; Howe and Jander, 2008; Ross, 1961). The best studied are the jasmonic acid (JA) and salicylic acid (SA) plant immune pathways. Broadly, against necrotrophs and piercing-sucking herbivores, they employ the JA pathway, which can produce proteinase inhibitors (PIs) and polyphenol oxidase (PPO) (Vijayan et al., 1998; Wang and Wu, 2013) while against biotrophic pathogens like viruses, they employ the SA pathway which produces pathogenesis-related-genes and systemic acquired resistance (Klessig and Malamy, 1994; Raskin, 1992). The SA and JA immune pathways are often reciprocally antagonistic meaning it is not always possible to mount an effective immune response against diverse pathogens (Gupta et al., 2000; Kunkel and Brooks, 2002; Thaler et al., 2002; Zarate et al., 2007). The beneficial effect of plant viruses on vector and non-vector fitness may occur via negative immune cross talk between the JA and SA pathways (Zhang et al., 2012). Plant viruses can activate the salicylic acid pathway (López-Gresa et al., 2016; Malamy et al., 1990; Singh et al., 2004) which can prevent activation of the JA pathway (Zhang et al., 2012). Another possibility is that facilitation is resource-based; viruses have been shown to increase the production of free amino acids in the plant host which may make it easier to feed upon (Blua et al., 1994; Nachappa et al., 2013; Shrestha et al., 2012).

In this study we investigate facilitation of non-vector macroparasites, the spider mites, *Tetranychus evansi and T. urticae*, by tomato spotted wilt virus (TSWV) in coinfections. Interactions between *Tetranychus spp.* and tomato immune responses have been well studied (Kant et al., 2004; Sarmento, Lemos, Bleeker, et al., 2011). Both T*. evansi* and *T. urticae* are negatively affected by the JA pathway via the production of proteinase inhibitors (Ament et al., 2004; Ataide et al., 2016). However, strategies in both exist to overcome JA defences. *T. evansi* can suppress or reduce induction of the JA pathway (Knegt et al., 2020; Sarmento, Lemos, Bleeker, et al., 2011; Teodoro-Paulo et al., 2023). *T. urticae* has generally been found to induce the JA pathway (Kant et al., 2004; Li et al., 2002; Rioja et al., 2017). However, this can be induced to relatively low levels (Teodoro-Paulo -preprint; (Kant et al., 2008), and some strains or populations are tolerant to elevated JA (Kant et al., 2008).

Previous work has shown that *T. urticae* have higher fecundity on TSWV infected host plants (Belliure et al., 2010; Nachappa et al., 2013). We were interested in the generality of facilitation of spider mites by TSWV, to better understand whether it occurs for different TSWV strains, different *Tetranychus* species and on different varieties of tomato plant. We first describe 2 experiments where we explore facilitation of *T. urticae* on tomato leaves cut from plants infected with TSWV, a commonly used method with the spider mite system (Teodoro-Paulo et al., 2023) as plant immune responses can still be induced in cut leaves (Dias et al., 2022; Stewart, 1990). We then describe experiments on whole plants in which we show a positive effect of TSWV on *T. evansi* and *T. urticae* on 2 different varieties of tomato plant. We find that different strains of TSWV can positively impact *T. urticae*, that both *T. evansi* and *T. urticae* benefit from facilitation and that it occurs on different varieties of tomato plant.

## Material and Methods

### Tomato spotted wilt virus

Tomato spotted wilt virus (TSWV) is a tripartite RNA plant virus from the Tospoviridae family with 2 ambisense RNA strands (Best, 1968; Van den Hurk et al., 1977). It is one of the most menacing plant viruses for agriculturists globally, and destroys ornamental plants, crops, and vegetables (Crosslin et al., 2009; Roselló et al., 1996). The symptoms include stunted growth in plants, leaf and stem necrosis, chlorotic spots and spoiled fruits (Roselló et al., 1996; Sherwood et al., 2003). In nature, TSWV is vectored by thrips, mainly, Frankiniella occcidentalis (Ullman et al., 2002; Whitfield et al., 2015), but in our laboratory, we mechanically inoculate the host plants with a virus inoculum prepared by using leaves of an already infected plant. The infected leaf (1 g) is crushed with 4 mL of buffer (Na_2_HPO_4_.12H_2_O 0.03 M + 0.2% DIECA) and 90 mg of charcoal in a pestle and mortar until it becomes a smooth paste. Then we add 75 mg of carborundum as a surface abrasive for the solution to be easily absorbed by the plant cells. This inoculum is rubbed gently with a finger on the surface of the 1st fully expanded leaf to coat the entire surface and rinsed after 20-25 minutes. The virus strains were obtained from the Plant Pathology unit, INRAE, Avignon and were maintained in the laboratory by serial passage of new plants every 14 days on 3-week-old tomato plants (variety Moneymaker). Plants are checked for infection using ELISA test kits by Agdia. TSWV infected plants are maintained at 25 ± 2°C with a 16: 8 light: dark cycle.

### The spider mites

Tetranychus urticae, is a generalist mite species feeding on >1000 species of agricultural plants (Helle and Sabelis, 1985; Shih et al., 1976) while Tetranychus evansi are specialists on the Solanaceae family (Qureshi et al., 1969). Spider mites are haplo-diploid. Females lay eggs which hatch ∼4 days later. The juvenile period includes 1 nymph stage and 2 deutonymph stages with adults emerging 13-14 days later. The populations of both species used in these experiments are outbred populations which originated and were created in Portugal (see (Godinho et al., 2020) for more details)). Subpopulations of each were transferred to Montpellier in January and November 2022, and have since been maintained on 3 Moneymaker tomato plants in big plastic boxes (dimension 520 mm x 300 mm x 250 mm) with one plant being changed every week. Populations are kept at 25 ± 2 C ° on a 16: 8 light: dark cycle. Prior to each experiment, groups of 50 mated females were placed together on a cut tomato leaf to lay eggs to produce equally aged daughters for the experiments.

### Host Tomato Plants

We used different commercially available varieties of host tomato plants, Solanum lycopersicum, in our experiments; Moneymaker, Saint-Pierre, Olympe and Microtom. Moneymaker and Olympe are F1 varieties. All plants were grown in an arthropod free environment at 25°C ± 3°C with a 16h:8h light: dark cycle.

## 1. Effect of tomato variety on the interaction between TSWV and *T. urticae* on isolated tomato leaves

### Experiment 1

This experiment tested the effect of TSWV on Tetranychus urticae life-history traits on 3 different varieties of tomato, Moneymaker (MM), Saint-Pierre (SP), and Olympe (OL). Eight plants of each variety were infected with the France81 strain of TSWV 3-weeks post sowing. Eight control plants of each variety were also mock infected by applying a mock inocula (buffer with uninfected plant leaves) to the first fully expanded leaf. Infections were confirmed 14 days later by removing a small piece of leaf from infected plants, and testing with TSWV ImmunoStrips (from Agdia company, Paris, France). Small pieces of leaflets were also removed from control plants to control for any effect of removing leaves when testing for infection.

We cut the 2 leaves above the inoculated one and put both into the same transparent, plastic box (250 mm x 180 mm x 75 mm). Both leaves were pressed onto the wet cotton roll, and the stem wrapped in wet cotton wool to keep them fresh. We placed 10 adult mated T. urticae females onto 2 leaflets on one of the leaves and 1 leaflet on the second leaf (both from the same plant, both the leaves in a box representing a single plant replicate) (Figure 1). A barrier of lanolin and tanglefoot glue mixed in a 1:1 ratio was applied to the stem to isolate the mites on a single leaflet. Fecundity (eggs laid by the females) was measured on the 4th day after transferring the mites to boxes at which point the mother females were removed. On the 14th day after transferring spider mites, we counted the total number of adult male and female offspring on the leaflets. The adult count was continued for 2 more days to account for temporal variation in development. We also measured the size of the leaflets by quantifying the area from photos taken on day 12-13 using Image-J. This experiment aimed to have 3 tomato varieties x 2 parasite treatments (1 virus + mock control) x 8 replicates = 48 plants (see Table S1 for actual replicate numbers).

**Figure 1:**
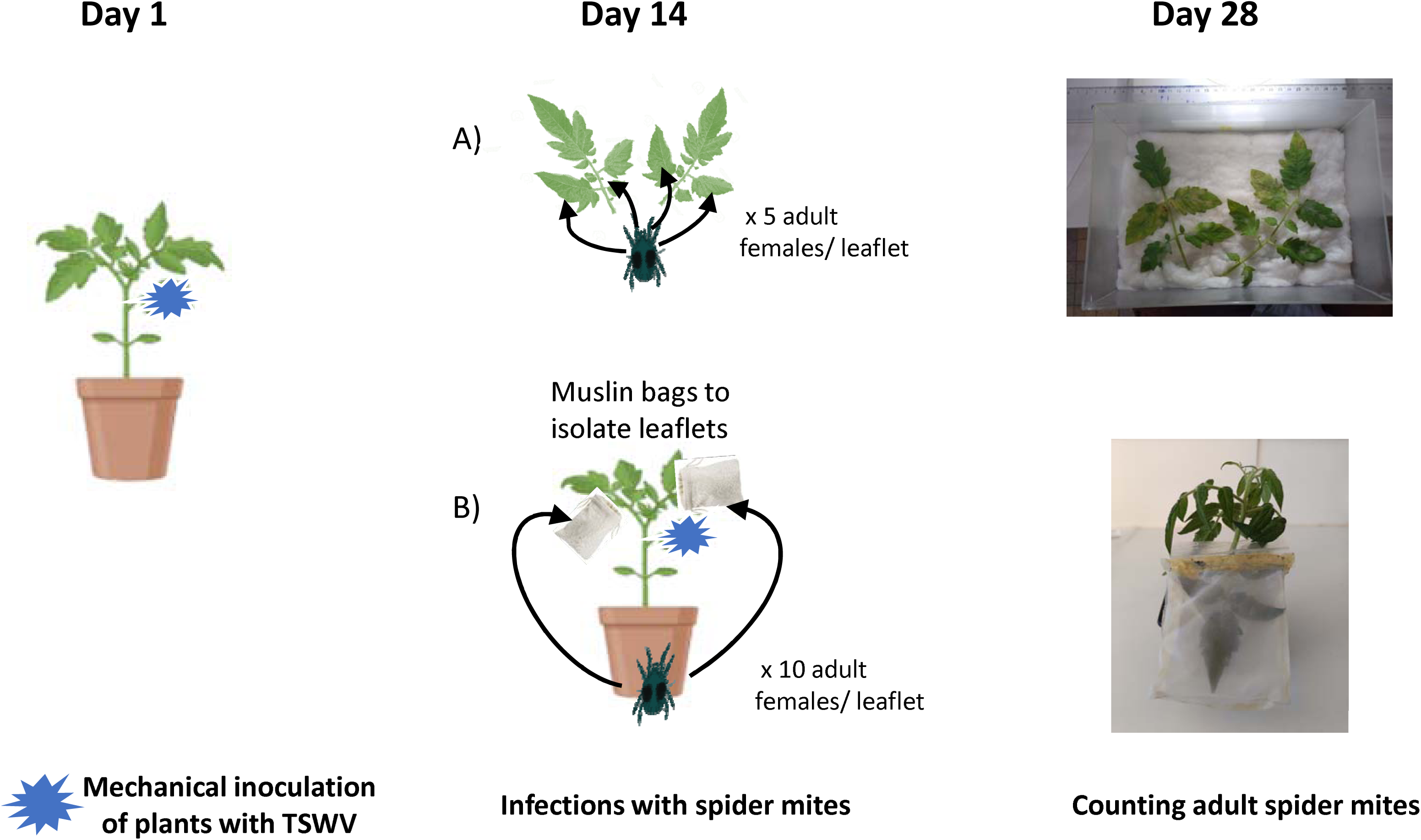
Pictorial representation of the experiment design on cut leaves and whole plants. A) spider mites transferred onto the cut leaves from the TSWV or mock infected plants used in experiments 1 and 2, while B) spider mites transferred onto the leaves of an intact plant used in experiments 3 and 4. In cut leaves, the spider mites were contained onto the leaflet with glue and tanglefoot as a barrier, while on the whole plants, we made muslin bags to isolate them onto the leaflet.

### Experiment 2

Here we tested the effect of three different strains of TSWV (France81, LYE1137vir and LYE55) on T. urticae life-history traits on Moneymaker leaves. As in Experiment 1, we infected 8 3-week old plants post sowing and had 4 plants that were mock inoculated as before. In addition here we also had 7 clean plants as controls to test for effects of mock inoculation. As before, 2 leaves were taken from each plant and placed on water saturated cotton wool in the same box and 10 female spider mites placed on 1 leaflet of each leaf isolated with lanolin and Tanglefoot glue (Figure 1). We counted the number of eggs 4 days later (and removed the mothers), the number of emerging juveniles on the 8th day, and the number of male and female adult offspring on the 14th day. We quantified the leaflet areas by taking photos and analysing them with Image-J. See Table S1 for final replicate numbers.

## 2. Effect of tomato variety on facilitation of *T. urticae* and *T. evansi* by TSWV on whole plants

### Experiment 3

In this experiment we tested the effect of TSWV on T. urticae and T. evansi life-history traits on whole tomato plants of the Moneymaker and Microtom varieties. We infected 16 plants of each variety 3 weeks post sowing with the France81 virus strain and mock inoculated another 16 as described above. After 14 days we removed a leaf from all plants, checked infected plants for virus infection and placed them in groups of 8 in plastic boxes (dimension 520 mm x 300 mm x 250 mm).

We transferred adult female mites of either T. urticae or T. evansi in groups of 10 onto one leaflet on two different leaves above the inoculated leaf on each plant. Here the mites were isolated onto their respective leaflet using a muslin leaf cage (dimensions 70mm x 90 mm x 10 mm). These cages were made by glueing muslin into a box shape and attaching a plastic ziplock from a freezer bag as an opening. The different treatments were randomised across boxes following the transfer of mites but mite species were kept in separate boxes due to contamination risks). We counted the number of adult male and female adults using a binocular microscope onthe 14th day following removal of the infested leaves. Eggs or other developmental stages prior to day 14 were not measured as it was not possible to remove and replace the muslin bag without disrupting the mites.

This experiment had 2 tomato varieties x 2 parasite treatments (1 virus + mock control) x 2 *Tetranychus* species x 8 replicates (see Table S1 for actual replicate numbers).

### Experiment 4

In a separate experiment, we tested whether TSWV infection influenced female survival and fecundity of *T. urticae* on Moneymaker plants. This was to be sure that differences in fitness on TSWV infected plants was not due to differential survival of mothers. We infected six plants with France81 and seven with LYE1137vir TSWV strains as described previously, and had six mock inoculated controls. After 14 days we checked the plants for positive infections and kept positive plants in plastic boxes (520 mm x 300 mm x 250 mm).

We transferred 10 adult female mites onto 1 leaflet each of 2 different leaves (on the 2 leaves above that which had been inoculated), and isolated them with the muslin leaf cages as described above. On the 4th day after transferring the mites, we took 3 plants that had been infected with LYE1137vir, 2 positively infected with France81, and 3 mock infected plants, and removed the leaves to count eggs; also, we counted the number of mother female mites surviving on the leaflet. The number of juveniles on the leaves on the 7 remaining plants (3 mock inoculated and 4 infected with LYE1137vir) were counted in the same way on day 8. This experiment had 3 parasite treatments (2 virus + mock control) x 6-7 replicates (see Table S1).

## Statistical analyses

All analyses were done in R version (4.3.2). All models were simplified in a stepwise fashion removing non-significant terms. Full models are shown in Tables in the Supplementary materials.

### Experiment 1

We used Generalized Linear Mixed Models (GLMMs) to test whether fecundity, total number of adult offspring, total number of adult daughters, and the offspring sex ratio were affected by plant variety, virus infection, leaflet area, and their interactions. Plant variety and virus infection were included in models as fixed factors and leaf area as a covariate. For fecundity (total number of eggs), total number of adults and the total number of females, GLMMs with a negative binomial distribution were implemented using the package glmmTMB as these count variables were overdispersed. Offspring sex ratio was analysed using a GLMM in glmmTMB with a binomial error structure and logit link function with the columns for the number of adult males and the total number of females combined as a response variable using cbind. Plant replicates nested within viral infection were included in all models as a random factor. Leaflet was included in the sex allocation model as an observation level random effect to control for overdispersion (Harrison, 2014).

### Experiment 2

We used GLMMs to test whether total fecundity, number of juveniles, number of adult offspring, number of adult daughters, and the offspring sex ratio were affected by viral strain, leaflet area, and their interaction. Viral strain was included in models as a fixed factor and leaflet area as a covariate. For fecundity, the number of juveniles, total adults, and number of female adults, GLMMs with a negative binomial distribution were implemented using the package glmmTMB. Offspring sex ratio was analysed using a GLMM with a binomial error structure and logit link function. Individual plant replicates nested within viral treatment were included in all models as a random factor. The models for fecundity and sex ratio also included leaflet as an observation level random effect to control for overdispersion (Harrison, 2014).

### Experiment 3

We used GLMMs to investigate how TSWV infection, mite species and plant variety influenced the total number of adults, the number of adult females and offspring sex ratio. Infection treatment, spider mite species and plant variety and their interactions were included in models as fixed effects. For the total number of adults and the total number of females, GLMMs with a negative binomial distribution were implemented; for the sex ratio model, a GLMM with a binomial error structure and logit link function was used. Individual plant replicate nested within box ID, and box position on the shelf were included in all models as a random factor. Leaflet was also included in the models for fecundity and sex allocation as an observation level random effect to control for overdispersion (Harrison, 2014).

### Experiment 4

We used GLMMs to investigate how TSWV infection affected the number of eggs, number of juveniles, and female survival on each leaf. Viral treatment was included in these models as a fixed factor with 3 levels (France81, LYE1137vir and mock). For the number of eggs and mothers surviving, we used a GLMM with Poisson error structure and log link function using the package glmmTMB. In a separate model, we investigated how TSWV infection (LYE1137vir or mock) affected the number of juveniles on each leaf using a negative binomial distribution with glmmTMB. Individual plant replicates nested within viral treatment were included in models as a random factor.

## Results

### Experiment 1: Effect of tomato variety on the interaction between TSWV and T. urticae on isolated tomato leaves

There was no main effect of TSWV infection on *T. urticae* fecundity (𝛘^2^_1_ = 0.18, p = 0.6706), the total number of adult offspring (𝛘^2^_1_ = 0.10, p = 0.756), or the number of adult daughters (𝛘^2^_1_ = 1.06, p = 0.303). TSWV infection increased the number of adult daughters (variety*virus infection 𝛘^2^_2_ = 11.06, p = 0.004) and total number of offspring becoming adult (variety*virus infection 𝛘^2^_2_ = 6.88, p = 0.032) on the Moneymaker tomato variety only as shown by significant interactions between virus infection and plant variety (Figure 2a; Table S2). There was also a significant interaction between leaf area and virus infection for fecundity (𝛘^2^_1_ = 6.27, p = 0.0123), number of adult daughters (𝛘^2^_1_ = 8.09, p = 0.004) and number of adult offspring (𝛘^2^_1_ = 10.85, p < 0.0001) showing a positive relationship among these traits on control, but not virus infected plants.

**Figure 2:**
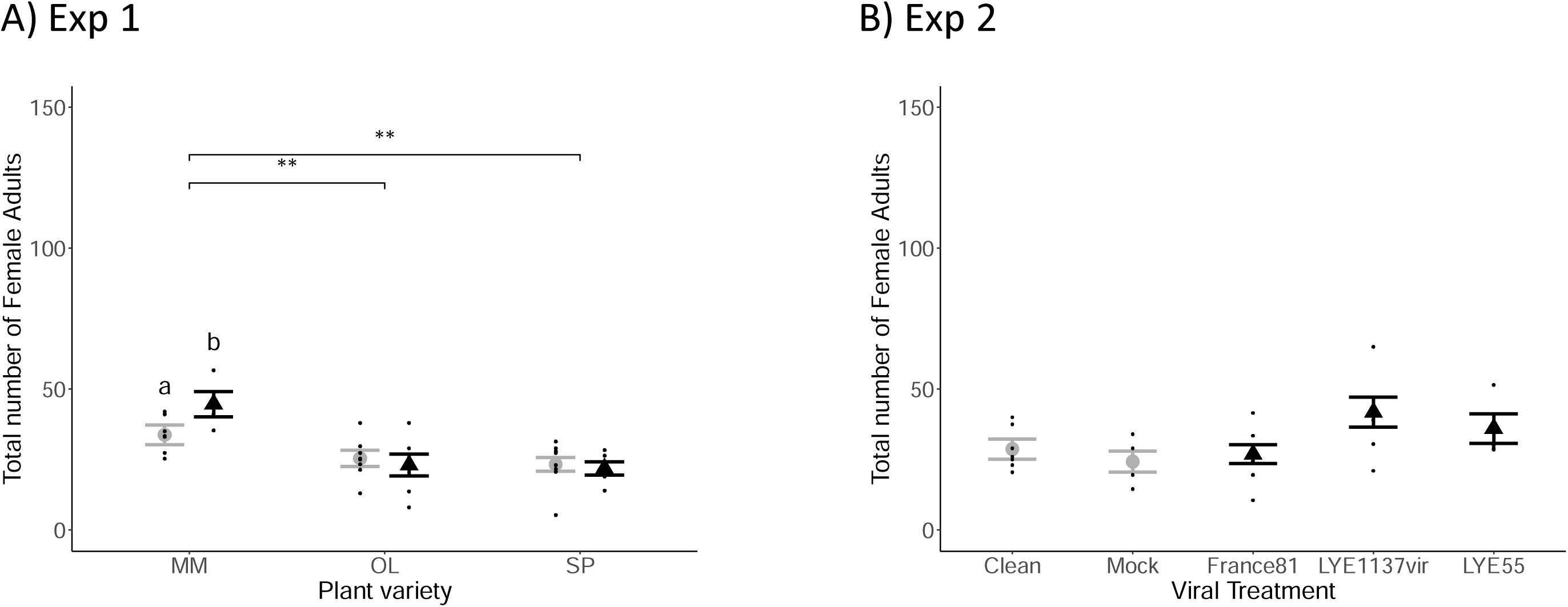
Effect of tomato spotted wilt virus on mean (± standard error) total number of adult female mites in A) Experiment 1, on mock (gray) and TSWV infected (black) plants of Moneymaker (MM), Olympe (OL) and Saint-Pierre (SP) tomato varieties. ** represent the significant differences among the plant varieties, while “a” and “b” represent the significant difference between the mock and virus infected in Moneymaker variety. B) Experiment 2, on mock and clean controls (gray) or plants infected with 3 different TSWV strains (France81, LYE1137vir, LYE55) (black). Mean is represented by a gray dot in all non-TSWV treatments, and by a black triangle in all TSWV treatments. All the data points are jittered in black along the respective treatments.

There was a significant effect of tomato variety showing higher *T. urticae* fecundity (𝛘^2^_2_ = 11.99, p = 0.0025), more adult offspring (𝛘^2^_2_ = 35.07, p < 0.001) and more adult daughters (𝛘^2^_2_ = 17.41, p < 0.001), on Moneymaker plants (Figures 2a and S1, Table S2). There was no effect of TSWV infection, tomato variety or leaflet area on the offspring sex ratio (Figure S2).

### Experiment 2

There was a marginally significant effect of viral strain with *T. urticae* having a higher number of adult daughters (𝛘^2^_4_ = 11.34, p = 0.023) on leaves from plants infected with the LYE1137vir and LYE55 viral strains compared to control plants (both clean and mock infected) and plants infected with the France81 viral strain (Figure 2b). There was also a more female biased offspring sex ratio (𝛘^2^_4_ = 22.03, p = 0.0002) on plants infected with the LYE1137vir strain. There was no effect of viral strain, leaflet area or their interaction on fecundity, the number of juveniles or the total number of adults (Figures S3 and S4, Table S3).

## Effect of TSWV infection on *T. urticae* and *T. evansi* life-history traits on whole plants

TSWV infection increased the number of offspring becoming adults (TSWV infection; 𝛘^2^_1_ = 16.29, p < 0.0001) and the number of adult daughters (TSWV infection; 𝛘^2^_1_ = 14.85, p < 0.0001). There were also more adult offspring (plant variety; 𝛘^2^_1_ = 4.22, p = 0.04) and adult daughters (plant variety; 𝛘^2^_1_ = 4.87, p = 0.027) on Moneymaker compared to Microtom plants. Overall *T. evansi* had a higher total number of adult offspring (mite species; 𝛘^2^_1_ = 5.38, p = 0.02) and more adult daughters (mite species; 𝛘^2^_1_ = 10.68, p = 0.001) than *T. urticae* on both plant varieties (see Table S4). There were no significant interactions among TSWV infection, mite species and plant variety on the number of adult offspring or daughters (Figure 3, Table S4).

**Figure 3:**
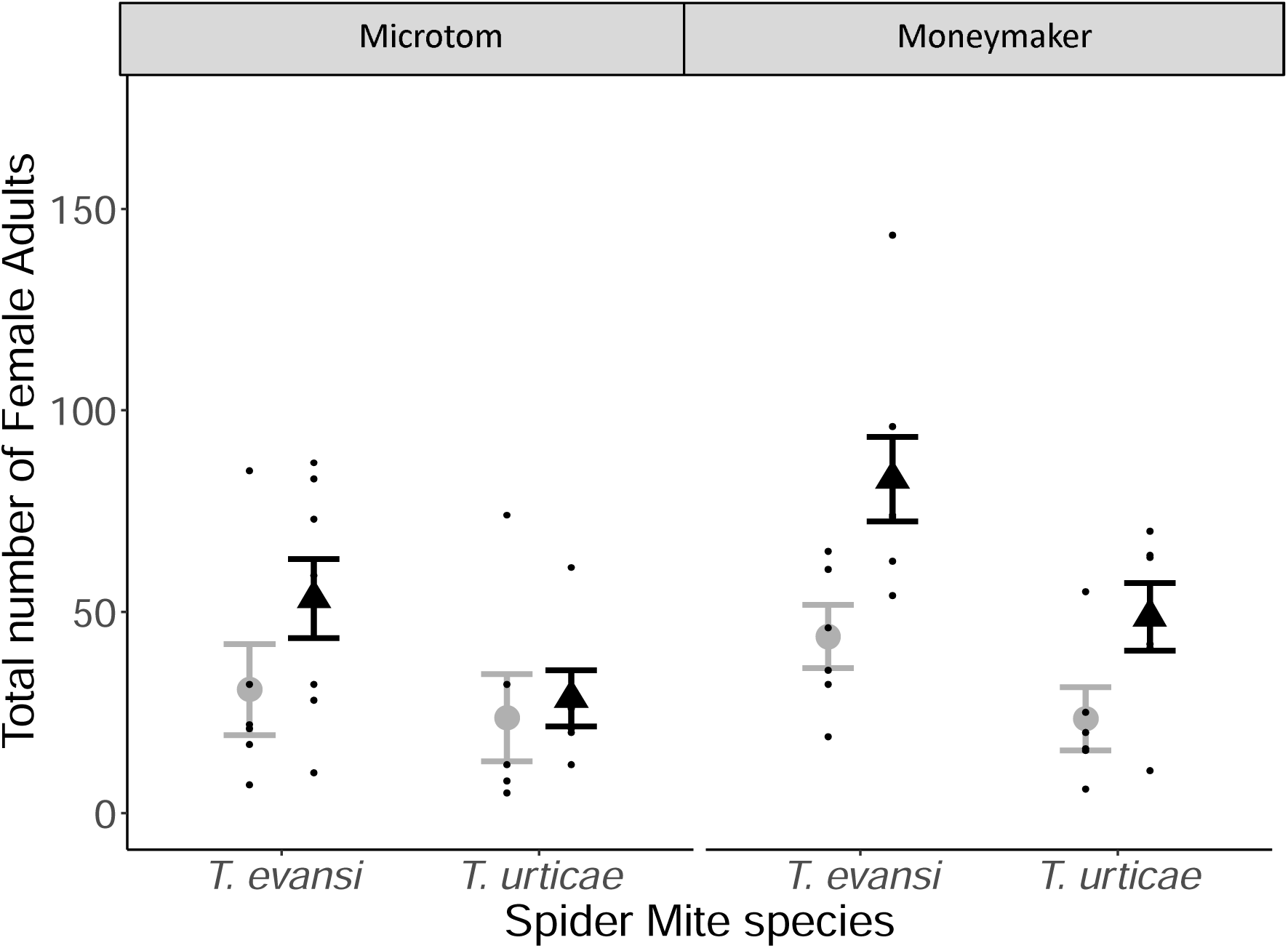
Effect of tomato spotted wilt virus on mean (± standard error) *Tetranychus urticae* and *Tetranychus evansi* number of adult daughters on mock (gray) and virus infected (black) plants of the Microtom and Moneymaker varieties. Mean is represented by a gray dot in all non-TSWV treatments, and by a black triangle in all TSWV treatments. All the data points are jittered in black along the respective treatments.

The effect of TSWV infection on offspring sex ratio changed according to plant variety (virus infection*plant variety; 𝛘^2^_1_ = 7.17, p = 0.0074), with a less female biased sex ratio on virus infected plants of the Microtom variety, and mite species (virus infection*mite species; 𝛘^2^_1_ = 10.1, p = 0.0015) with *T. urticae* having a more female biased offspring sex ratio on virus infected plants (see Figure S5).

There was a significant effect of TSWV infection on *T. urticae* fecundity (𝛘^2^_1_ = 5.938, p = 0.0513) with more eggs being laid on virus infected plants. Similarly, the number of eggs that hatched, measured as the number of juveniles was higher in the presence of the virus (𝛘^2^_1_ = 7.312, p = 0.0068; see Figure S6). There was no difference in the survival of mothers on virus infected or control plants (𝛘^2^_1_ = 0.056, p = 0.97212) (Figure 4; Table S4) meaning that there was no role of differential survival of mothers in facilitation.

## Discussion

Our results confirm the facilitatory effect of Tomato spotted wilt virus on both *T. urticae* and *T. evansi* on whole plants. This advances on previous studies (Belliure et al., 2010; Nachappa et al., 2013) showing that TSWV has a positive effect on the number of adult daughters and total number of adult offspring in addition to fecundity. Facilitation was repeatable for both *T. evansi* and *T. urticae* on two different tomato varieties on whole plants, Moneymaker and Microtom. In contrast, on cut leaves, facilitation was variable across experiments and depended upon the tomato variety. We attribute this to the non-homogeneous spread of virus throughout the plant meaning the cut leaves used in the experiment were not always infected with TSWV or an indirect systemic effect mediated via virus infection was not effective.

**Figure 4:**
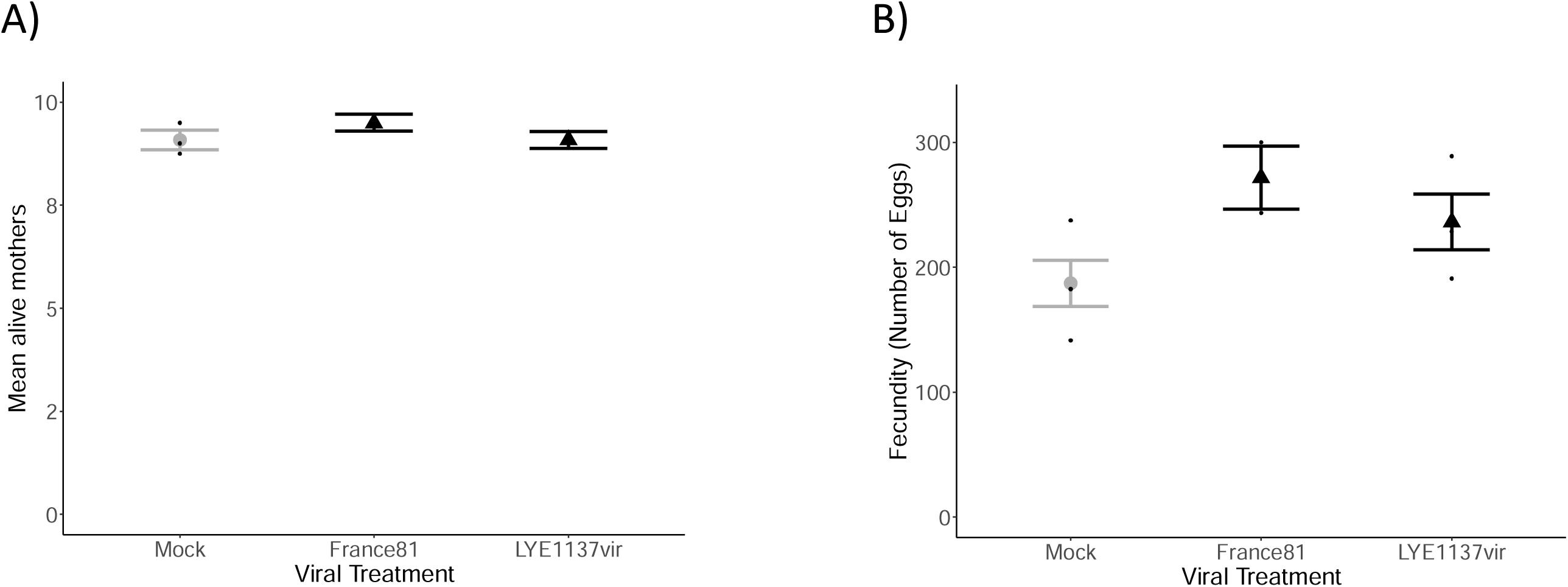
Effect of tomato spotted wilt virus on mean (± standard error) *T. urticae* A) mother survival, B) fecundity on Moneymaker whole plants mock infected (gray) or infected with TSWV strains France81 and LYE1137vir (black). Mean is represented by a gray dot in all non-TSWV treatments, and by a black triangle in all TSWV treatments. All the data points are jittered in black along the respective treatments.

### Facilitation on whole plants

The mechanism of facilitation of *T. urticae* and *T. evansi* by TSWV might be due to different, non-mutually exclusive, mechanisms. A number of studies show that facilitation of arthropods (vectors and non-vectors) by plant virus infection arises via negative immune cross talk between the JA and SA immune pathways (Shi et al., 2014; Zhang et al., 2012). Prior induction of the SA pathway following infection with TSWV (López-Gresa et al., 2016) may prevent upregulation of the JA pathway and production of proteinase inhibitors which disrupt spider mite feeding and digestion of the plant material (Sarmento, Lemos, Bleeker, et al., 2011).

As *T. evansi* have been shown to suppress the JA pathway or induce it to low levels (Knegt et al., 2020; Sarmento, Lemos, Bleeker, et al., 2011; Teodoro-Paulo et al., 2023), we did not think that they would necessarily benefit from TSWV facilitation, if mediated via negative immune cross talk (Thaler et al., 2010; Zhang et al., 2012). However, JA can negatively impact *T. evansi* fecundity at very low concentrations (Ataide et al., 2016). As the population used in this experiment does not suppress JA to levels lower than in clean plants (Teodoro-Paulo et al., 2023) they may benefit from any prevention of JA expression.

Another, non-mutually exclusive hypothesis is that facilitation is via TSWV changing the nutritional quality of the plant. Plant viruses, including TSWV, can increase levels of free-amino acids circulating in cell sap with positive effects on arthropod life-history traits (Ángeles-López et al., 2018; Blua et al., 1994; Nachappa et al., 2013). Thus, increased free-amino acids may be a source of nitrogen or essential amino acids for both *Tetranychus* species. Finally, the virus capsid itself may be a source of amino acids for the spider mites as they possibly ingest some of the virus particles. The capsid is mainly composed of nucleoproteins arranged in spherical or icosahedral form. A study by (Orlob, 1968) shows that *T. urticae* can ingest different viruses that can also be detected in the excretory material.

### No facilitation on cut leaves

All plants used in our experiments had tested positive for TSWV infection (for plants in virus treatments). However, despite plants being infected with TSWV, facilitation of spider mites was not always observed when measured on cut leaves. This may be for a number of reasons.

One possibility is non-homogeneous spread of virus across the plant, and thus, leaves upon which facilitation was not observed may have been uninfected. Viruses are only found in leaves that mature following infection (Agrios, 2005). This is because viruses exploit existing source to sink transport routes in the plant for solutes (Carrington et al., 1996). The source-to-sink transport system describes how mature ‘source’ leaves produce sugars (and other essential nutrients) which are transported to younger ‘sink’ leaves where they are used (Agrios, 2005; Lemoine et al., 2013)). Following inoculation, the virus spreads from the epidermal cells via cell-to-cell movement through the plasmodesmata to the phloem in the sieve elements to be transported to the roots and other sink leaves (Hipper et al., 2013). (Samuel, 1934) described how tobacco mosaic virus in potato plants first travels to the roots, then the apical leaves and then gradually down the plant. Consistent with this we found that apical tomato leaves were always infected 11 days following infection but that leaves immediately below were not always infected (see figure S7). Leaves below became infected sequentially from top to bottom (Supplementary material A1). This means that leaves used in our experiments will not always have contained viral particles which may have prevented facilitation occurring. However, we did not think that removing leaves would necessarily prevent facilitation as plant mediated effects were maintained on cut leaves in other experiments with spider mites and other arthropods (Dias et al., 2022; Musser et al., 2002; Sarmento, Lemos, Dias, et al., 2011). For example, induction of plant immune responses on whole plants by *T. urticae* can later reduce *T. evansi* fecundity on leaves or leaf discs that have been removed from the plant, including on leaves never previously in contact with mites (Dias et al., 2022; Sarmento, Lemos, Dias, et al., 2011). Although the mechanism of facilitation is linked to systemic responses, variation in expression of genes implicated in the JA and SA pathways can be observed locally in different regions of the same leaf (Betsuyaku et al., 2018; Schimmel et al., 2017). Furthermore, levels of free-amino acids may be higher in leaves infected with TSWV as found by (Nachappa et al., 2013). Thus, facilitation of spider mites by TSWV may rely on local interactions. In line with this, it would be interesting to see if the degree of facilitation on whole plants depends on whether the leaf is infected with virus and viral quantity on whole plants. Other systems have found variation in effects of plant parasites/symbionts on arthropod fitness when measuring traits on whole plants versus leaf discs (Jaber and Vidal, 2010).

### Sex allocation in response to virus infection

We find that virus infection can change the offspring sex ratio, but not always in the same direction. *T. urticae* had a less female biased offspring sex ratio when on whole plants infected with TSWV but a more female biased sex ratio when on TSWV infected cut leaves, compared to uninfected plants or leaves. In contrast, *T. evansi* had a more female biased sex ratio when on whole plants infected with TSWV. The reasons for these differences in sex allocation, the differential investment into male versus female offspring, for *T. urticae* on whole plant versus cut leaves, and between the two species, are not clear. One possibility is that the effect of TSWV is related to resource availability. In solitary parasitoids (Charnov et al., 1981) and sawflies (Craig et al., 1992) allocation to female offspring is higher in good resource environments attributed to greater fitness returns from larger daughters and/or females having greater benefit from good environments. Although female spider mites are larger than males, it is not clear whether relative fitness returns are greater from larger daughters (Macke et al., 2010).

Fitness returns may arise due to the optimal offspring sex ratio which, in spider mites, will also ultimately result from higher allocation of resources to daughters. Spider mites are haplodiploid, females developing from fertilised eggs and males from unfertilised eggs. Previous work has shown that one way female *T. urticae* adjust their sex allocation is through allocation of resources to eggs, with larger eggs more likely to encounter sperm, be fertilised and become female (Macke et al., 2010). Further, females can facultatively adjust egg size and sex allocation adaptively in response to environmental cues (Macke et al., 2012). Thus, variation in sex allocation in the different environments may be due to resources available to allocate to eggs and offspring, with more females being produced in good resource conditions.

At the same time, overall we observe more female biased offspring sex ratios on leaves with higher densities. As females are the dispersing stage it has previously been postulated that *T. urticae* produce a more female biased offspring sex ratio under poor conditions to escape (Young et al., 1986). Density dependent dispersal has been shown for many species (Dorken and Pannell, 2008; Matthysen, 2005). Recently Chokechaipaisarn and Gardner (Chokechaipaisarn and Gardner, 2022) theoretically showed that more female biased offspring sex-ratios are favored under density-dependent dispersal in viscous populations, like spider mites.

### Virus interactions with non-vector species

Virus-vector interactions are indeed embedded within complex communities and better understanding of viral transmission may ultimately depend on knowledge about interactions at broader scales (Crowder et al., 2019; Eigenbrode et al., 2018). Viruses positively impacting the fitness of their vectors is adaptive if it increases transmission to new hosts (Blanc and Michalakis, 2016). However, it is not clear whether simultaneous facilitation of other non-vector parasites or arthropod herbivores sharing a host is costly which may in turn counter any beneficial effects on vectors. We are only aware of one study exploring the tripartite interaction between *T. urticae*, TSWV and their thrips vector and this study focussed on the positive effects of TSWV and thrips feeding on *T. urticae* life-history traits (Belliure et al., 2010). In the absence of virus, it has been shown that thrips predate upon *Tetranychus* species (Agrawal et al., 1999) with a higher proportion of *Frankliniella occidentalis* thrips larvae becoming adult when feeding on *T. urticae* eggs but a lower proportion becoming adult when feeding on *T. evansi* eggs (Ataide et al., 2019). Further, upregulation of the plant immune system by *T. turkestani* can negatively affect thrips survival in the absence of predation (Agrawal et al., 1999). These studies (as well as others) show that spider mites have the potential to impact thrips life-history via multiple mechanisms, notably, predation, immune mediated effects on the plant (Agrawal et al., 1999; Ataide et al., 2019), spider mite web protecting thrips from predation (Pallini et al., 1998) and possibly competition, but thrips were found to be the superior competitor (Zhi et al., 2006). Thus, overall it would seem that the presence of *Tetranychus* species positively impacts thrips life-history, and so would not necessarily affect TSWV transmission. Further work is needed to understand the direct effects of spider mites on TSWV growth and the potential for indirect effects via the thrips vector.

Greater knowledge about how virus - vector - non-vector interactions affect virus (or parasite) transmission and vector/non-vector life-history could aid control programs. In plant systems, vectors, non-vectors and viruses/parasites are often all pest species. It may be the case that one of the players disproportionately augments life-history traits and/or spread of others in the system. In such a scenario, targeting that species may have beneficial knock-on effects for the control of other pest species.

### More general evolutionary consequences of facilitation

Facilitation can be an important process shaping the ecology and evolution of both free-living and parasitic species (Bronstein, 2009; McIntire and Fajardo, 2014; Zélé et al., 2018). However, the potential for facilitation to persist and how evolution of facilitation proceeds between different species may depend on the consequences of facilitation for the benefactor (Bronstein, 2009) which may vary across environments. For example, facilitation is often predicted to be more prevalent under stressful conditions when competitive interactions are reduced (He et al., 2013; Hesse et al., 2021). In many contexts, including between plant viruses and their vectors and non-vectors, facilitation is a public good with the potential to benefit many organisms in the community. The production of metal detoxifying siderophores by some bacteria can facilitate others in a community without necessarily negatively affecting the siderophore producers, possibly due to localised facilitation or reduced competition in stressful environments (Hesse et al., 2021). However, co-evolution in coinfection in a nematode host between the protective bacteria *Enterococcus faecalis* and *Staphyloccucus aureus* selected against siderophore production by the latter; siderophores positively impact *E. faecalis* growth with negative effects on *S. aureus* (Ford et al., 2016). This highlights that facilitation can be selected against when pairwise interactions are at play. It may though be the case that scaling up to multiple interactions in a community may be more likely to permit niche differentiation and/or indirect interactions among players to maintain facilitative interactions.

Other studies have explored how vector - non-vector interactions may impact viral transmission. One study found that feeding by a non-vector species displaced an aphid vector to the most susceptible region of the plant for Pea enation mosaic virus transmission (Chisholm et al, 2019), and that the presence of those non-vector weevils increases the PEMV titre (Chisholm et al., 2018). (Su et al., 2016) found that non-vector *T. urticae* negatively affects the whitefly vector life-history traits on tomato hosts both in the presence and absence of Tomato yellow leaf curl virus (TYLCV). However, a more recent study by the same authors found that *T. urticae* actually increases TYLCV viral titre and also transmission by increasing whitefly feeding rate (Su et al., 2020). Consistent with this, a theoretical model shows that interspecific interactions with competitors, mutualists or predators affecting vector feeding or host choice impacted viral transmission more than when vector traits such as birth or death were affected (Crowder et al., 2019). These studies suggest that plant viruses can under certain circumstances actually benefit from interactions between their vectors and non-vector species.

## Conclusion

We find that TSWV can facilitate life-history traits of both *T. urticae* and *T. evansi* on whole plants. The reciprocal effect of spider mites on TSWV either via the plant or vector remains to be shown. Nevertheless, this work highlights that facilitation among parasites may be an important factor affecting parasite severity and spread. This work shows the importance of garnering a better understanding of parasite life-history within a multiple-parasite/community context.

## Data availability

Data will be made available in DRYAD.

## Acknowledgements

We most sincerely thank Dr. Benoit Moury and Marion Szadkowski from Plant Pathology unit, INRAE, Avignon, for providing our laboratory with the TSWV strains, and their helpful feedback about the experiments and virus pathology. We would also like to thank Dr. Yannis Michalakis for the insightful discussions about the results, and the project. We would like to thank Giacomo Zilio for helpful comments on a previous version of this manuscript.

