## Supplementary material for "Tomato spotted wilt virus facilitates non-vector spider mite species (*Tetranychus urticae* and *Tetranychus evansi*) on whole tomato plants": Table S1, Table S2, Table S3, Table S4, Figure S1, Figure S2, Figure S3, Figure S4, Figure S5, Figure S6, Figure S7, Supplementary material A1

### Supplementary Materials

Table S1: Total number of final replicates per treatment (numerator) over the initial number of replicates (denominator). Note that lower numbers were due to plants not testing positive for virus infection.

| | **Experiment 1** | | | | | | **Experiment 2** | | | | | **Experiment 3** | | | | | | | | **Experiment 4** | | |
|---|---|---|---|---|---|---|---|---|---|---|---|---|---|---|---|---|---|---|---|---|---|---|
| Spider mites | *T. urticae* | | | | | | *T. urticae* | | | | | *T. urticae* | | | | *T. evansi* | | | | *T. urticae* | | |
| Plant variety | Moneymaker | | Olympe | | Saint-Pierre | | Moneymaker | | | | | Moneymaker | | Microtom | | Moneymaker | | Microtom | | Moneymaker | | |
| Virus treatment | Control | Strain France 81 | Control | Strain France81 | Control | Strain France81 | Clean plants | Control | France81 | LYE55 | LYE1137vir | Control | France81 | Control | France81 | Control | France81 | Control | France81 | Control | France81 | LYE1137vir |
| Replicates | 8/8 | 5/8 | 8/8 | 5/8 | 8/8 | 6/8 | 7/7 | 4/4 | 7/8 | 5/8 | 5/8 | 6/6 | 7/8 | 7/7 | 8/8 | 6/6 | 7/8 | 7/7 | 8/8 | 6/6 | 2/8 | 7/8 |

Table S2: Output of GLMMs investigating the effect of tomato variety, TSWV infection and leaftlet area on *Tetranychus urticae* fecundity, number of offspring becoming adult, number of adult daughters, and the offspring sex ratio. The error distribution attributed in each model is stated in brackets after the variable name. The results shown (dF = degrees of freedom, Chisq = the chi square value, and p-value) are from the minimal model containing each term. Signiifcant terms are shown in bold. The variance components (± standard error) are shown for the random terms.

| | **Fecundity (nbinom)** | | | **No of adults (nbinom)** | | | **No of females (nbinom)** | | | **Sex ratio( binom)** | | |
|---|---|---|---|---|---|---|---|---|---|---|---|---|
| | dF | Chisq | p-value | dF | Chisq | p-value | dF | Chisq | p-value | dF | Chisq | P-value |
| **Variety** | **2** | **11.99** | **0.0025** | **2** | **35.07** | **<0.001** | **2** | **17.41** | **0.0002 *** ** | 2 | 1.54 | 0.4624 |
| **Virus infection** | 1 | 0.18 | 0.6706 | 1 | 0.097 | 0.7559 | 1 | 1.06 | 0.303 | 1 | 0.65 | 0.4212 |
| **Leaflet area** | 1 | 2.29 | 0.1304 | **1** | **12.98** | **0.0003 *** ** | **1** | **4.69** | **0.0302 *** | 1 | 0.22 | 0.6401 |
| **Variety: Virus infection** | 2 | 1.30 | 0.5214 | **2** | **6.88** | **0.0321 *** | **2** | **11.06** | **0.004 *** | 2 | 2.31 | 0.3151 |
| **Variety: Leaflet area** | 2 | 2.71 | 0.2576 | **2** | **5.81** | **0.0549 .** | **2** | **5.32** | **0.0698 .** | 2 | 3.89 | 0.1426 |
| **Virus infection: Leaflet area** | **1** | **6.28** | **0.0123** | **1** | **10.84** | **0.001 *** ** | **1** | **8.09** | **0.0045 ** ** | 1 | 0.24 | 0.6248 |
| **Variety: Virus infection: Leaflet area** | 2 | 2.02 | 0.3642 | 2 | 2.19 | 0.335 | 2 | 4.68 | 0.0965 . | 2 | 4.27 | 0.1182 |
| **Random effects** Replicate:Virus infection | 0.027 ± 0.16 | | | 0.01 ± 0.1 | | | 5.16e-10 ± 2.27e-05 | | | 3.74e-09 ± 6.11e-05 | | |
| Virus infection | 1.51e-10 ± 1.23e-05 | | | 1.84e-10 ± 1.36e-05 | | | 5.27e-11 ± 7.26e-06 | | | 5.29e-16 ± 2.29e-08 | | |
| **ID (each data point)** | | | | | | | | | | 0.49 ± 0.7 | | |

Table S3: GLMMs investigating the effect of different TSWV strains and leaflet area on *Tetranychus urticae* fecundity, number of offspring becoming adult, number of adult daughters, and the offspring sex ratio. The error distribution attributed in each model is stated in brackets after the variable name. The results shown (dF = degrees of freedom, Chisq = the chi square value, and p-value) are from the minimal model containing each term. Signiifcant terms are shown in bold. The variance components (± standard error) are shown for the random terms.

| | Fecundity (poisson) | | | Juveniles (nbinom) | | | No of adults (nbinom) | | | No of females (nbinom) | | | Sex ratio (binomial) | | |
|---|---|---|---|---|---|---|---|---|---|---|---|---|---|---|---|
| | dF | Chisq | p-value | dF | Chisq | p-value | dF | Chisq | p-value | dF | Chisq | p-value | dF | Chisq | p-value |
| **Virus infection** | 4 | 7.60 | 0.1073 | 4 | 5.25 | 0.2626 | 4 | 4.17 | 0.3837 | **4** | **11.34** | **0.023 *** | **4** | **22.03** | **0.0002 ***** |
| **Leaflet area** | 1 | 0.004 | 0.9511 | 1 | 0.08 | 0.7744 | 1 | 2.42 | 0.1198 | 1 | 1.96 | 0.162 | 1 | 1.75 | 0.186 |
| **Virus infection: Leaflet area** | 4 | 2.88 | 0.5778 | 4 | 4.06 | 0.3985 | 4 | 3.17 | 0.5306 | 4 | 4.39 | 0.3551 | 4 | 10.07 | 0.039 * |
| **Random effects** Replicate: Virus infection | 1.4e-09 ± 3.74e-05 | | | 2.78e-19 ± 5.28e-10 | | | 1.94e-09 ± 4.40e-05 | | | 2.11e-09 ± 4.59e-05 | | | 2.41e-02 ± 1.55e-01 | | |
| Virus infection | 2.27e-25 ± 4.77e-13 | | | 1.61e-10 ± 1.27e-05 | | | 3.44e-16 ± 1.85e-08 | | | 5.39e-15 ± 7.34e-08 | | | 1.21e-10 ± 1.1e-05 | | |
| **ID** | 0.11 ± 0.33 | | | | | | | | | | | | 0.085 ± 0.29 | | |

Table S4: GLMMs investigating the effects of different mite species, tomato variety and virus infection on the number of offspring bceoming adults, female adult offspring, and the offspring sex ratio. The error distribution attributed in each model is stated in brackets after the variable name. The results shown (dF = degrees of freedom, Chisq = the chi square value, and p-value) are from the minimal model containing each term. Signiifcant terms are shown in bold. The variance components (± standard error) are shown for the random terms.

| | **Experiment 3** | | | | | | | | | **Experiment 4** | | | | | | | | |
|---|---|---|---|---|---|---|---|---|---|---|---|---|---|---|---|---|---|---|
| | Mean alive mothers (poisson) | | | **No of eggs per female (poisson)** | | | **No of juveniles** | | | **No of adults (neg. binomial)** | | | **No of females (nbinom)** | | | **Sex ratio (binomial)** | | |
| | dF | Chisq | p-value | dF | Chisq | p-value | dF | Chisq | p-value | dF | Chisq | p-value | dF | Chisq | p-value | dF | Chisq | p-value |
| **Virus infection** | 2 | 0.06 | 0.9721 | **2** | **5.94** | **0.0513 .** | **1** | **7.31** | **0.0069 **** | **1** | **16.29** | **<0.001** | **1** | **14.85** | **0.0001 ***** | **1** | **0.01** | **0.922** |
| **Mite species** | | | | | | | | | | **1** | **5.38** | **0.0203 *** | **1** | **10.68** | **0.0011 **** | **1** | **15.01** | **0.0001 ***** |
| **Variety** | | | | | | | | | | **1** | **4.22** | **0.0398 *** | **1** | **4.87** | **0.0273 *** | **1** | **0.49** | **0.4845** |
| **Virus infection: Mite** | | | | | | | | | | 1 | 0.33 | 0.5665 | 1 | 0.06 | 0.8102 | **1** | **10.1** | **0.0015 **** |
| **Variety: Virus infection** | | | | | | | | | | 1 | 0.02 | 0.8791 | 1 | 0.55 | 0.4582 | **1** | **7.17** | **0.0074 **** |
| **Mite: Variety** | | | | | | | | | | 1 | 0.42 | 0.5172 | 1 | 0.29 | 0.586 | 1 | 0.34 | 0.5604 |
| **Virus infection: Mite: Variety** | | | | | | | | | | 1 | 0.15 | 0.6978 | 1 | 0.045 | 0.831 | 1 | 0.59 | 0.4399 |
| **Random effects** | | | | | | | | | | | | | | | | | | |
| Replicate: Virus infection | 5.78e-11 ± 7.60e-06 | | | 4.56e-03 ± 6.76e-02 | | | 2.56e-10 ± 1.6e-05 | | | | | | | | | | | |
| Virus infection | 1.44e-21 ± 3.79e-11 | | | 3.77e-11 ± 6.14e-06 | | | 1.59e-18 ± 1.26e-09 | | | | | | | | | | | |
| Replicate: Box | | | | | | | | | | 6.37e-02 ± 2.52e-01 | | | 2.76e-02 ± 1.66e-01 | | | 4.51e-10 ± 2.12e-05 | | |
| Box | | | | | | | | | | 9.53e-10 ± 3.09e-05 | | | 1.64e-09 ± 4.05e-05 | | | 1.98e-10 ± 1.41e-05 | | |
| Box.position | | | | | | | | | | 5.52e-03 ± 7.43e-02 | | | 1.60e-02 ± 1.26e-01 | | | 1.18e-02 ± 1.09e-01 | | |
| **ID** | | | | | | | | | | | | | | | | 2.44e-01 ± 4.94e-01 | | |

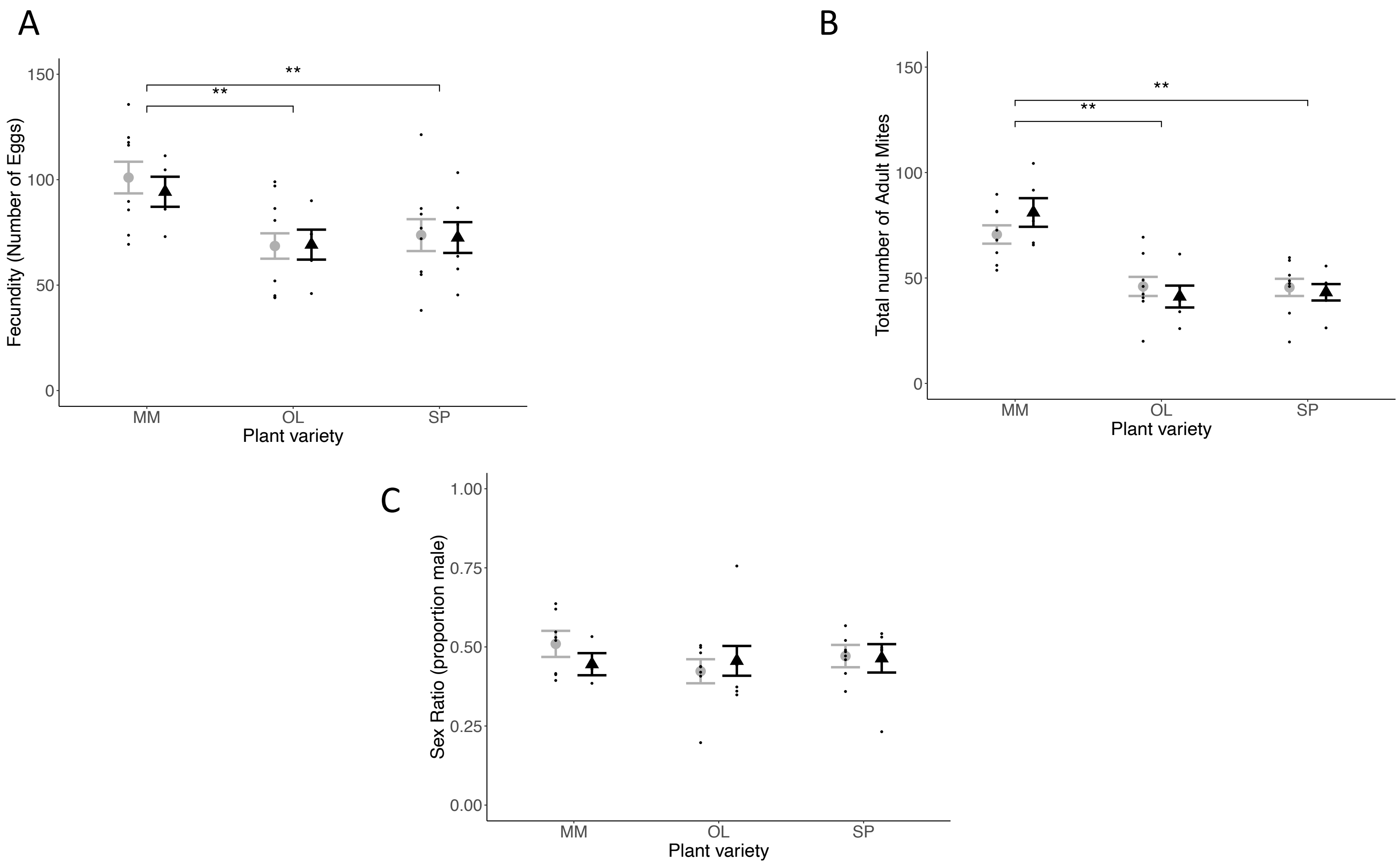


Figure S1: Effect of Tomato spotted wilt virus on mean (± standard error) A) fecundity (number of eggs laid) B) total number of adult mites, C) offspring sex ratio (proportion male) on mock (gray) and TSWV infected (black) plants of Moneymaker (MM), Olympe (OL) and Saint-Pierre (SP) tomato varieties. Mean is represented by a gray dot in all non-TSWV treatments, and by a black triangle in all TSWV treatments. All the data points are jittered in black along the respective treatments.

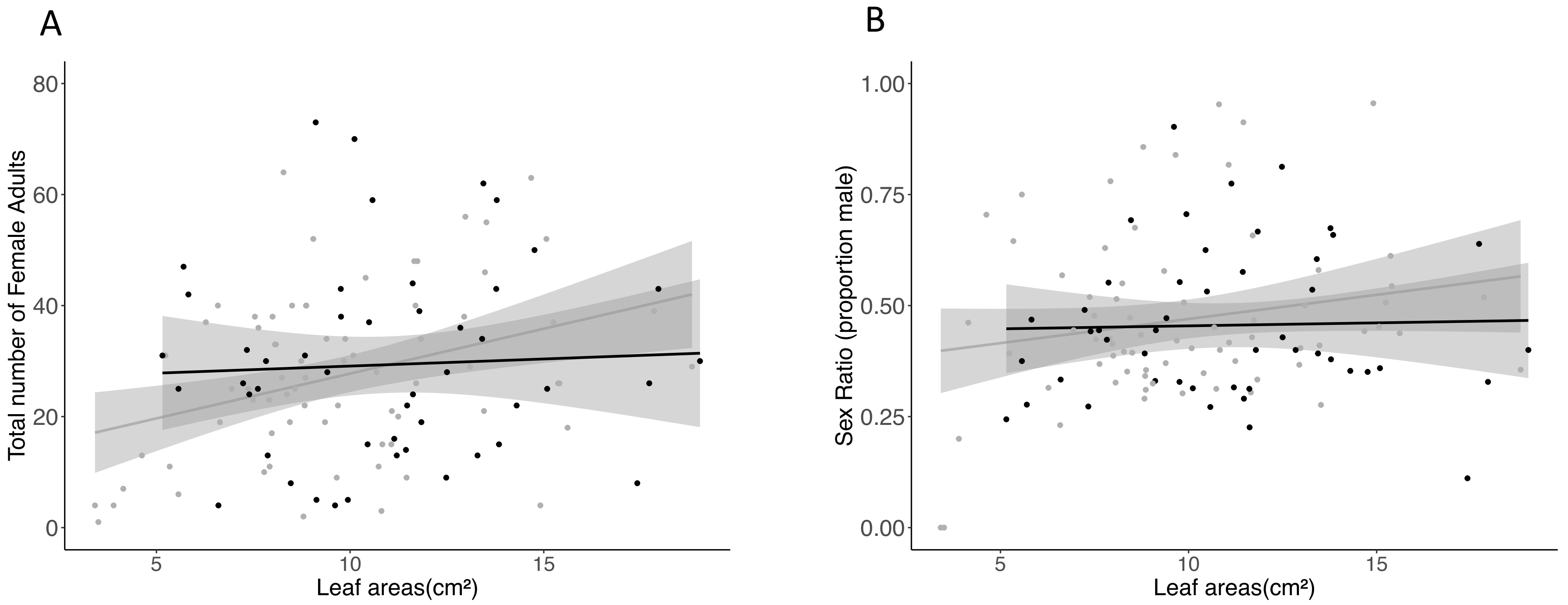


Figure S2: Effect of leaflet areas on A) total number of female adults, B) offspring sex ratio (proportion male) on mock (gray) and TSWV infected (black) leaflets of Moneymaker tomato plants (including leaflets of all three different varieties). Data points in black or gray belong to TSWV infected and mock respectively.

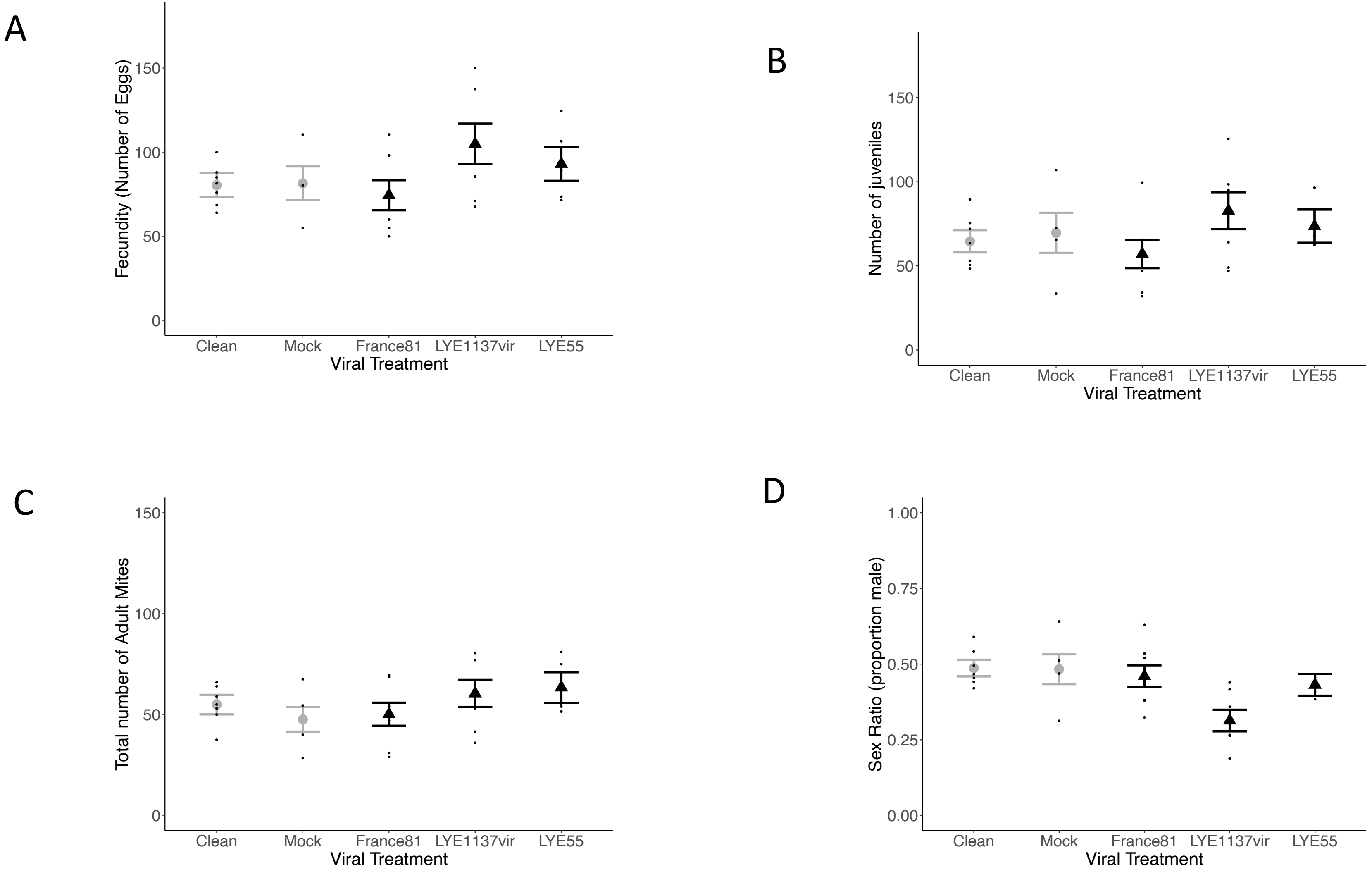


Figure S3: Effect of Tomato spotted Wilt Virus on mean (± standard error) A) fecundity (number of eggs laid) B) number of juveniles C) total number of adult mites, D) offspring sex ratio (proportion male) on mock or control (gray) or plants infected with 3 different TSWV strains (France 81, LYE 1137, LYE 55) (black). Mean is represented by a gray dot in all non-TSWV treatments, and by a black triangle in all TSWV treatments. All the data points are jittered in black along the respective treatments.

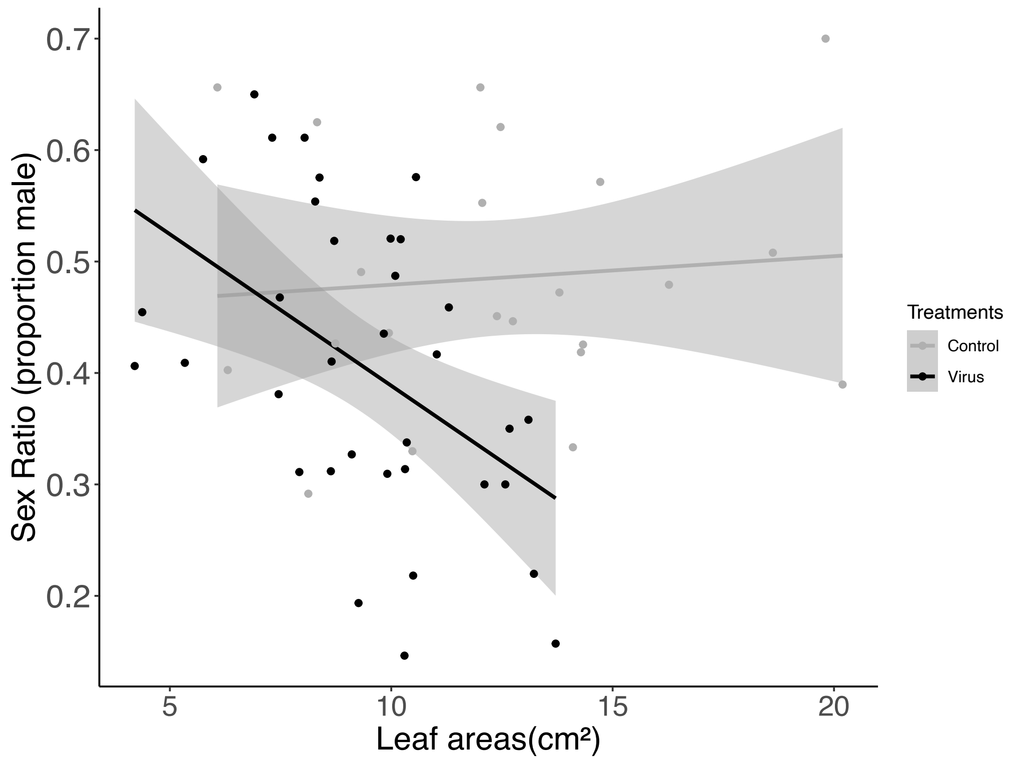


Figure S4: Effect of leaflet areas on offspring sex ratio (proportion male) on control (gray) and TSWV infected (black) leaflets of Moneymaker tomato plants (including leaflets of mock and clean in control treatment, and three different TSWV strains in Virus treatment). Data points in black or gray belong to TSWV infected and mock respectively.

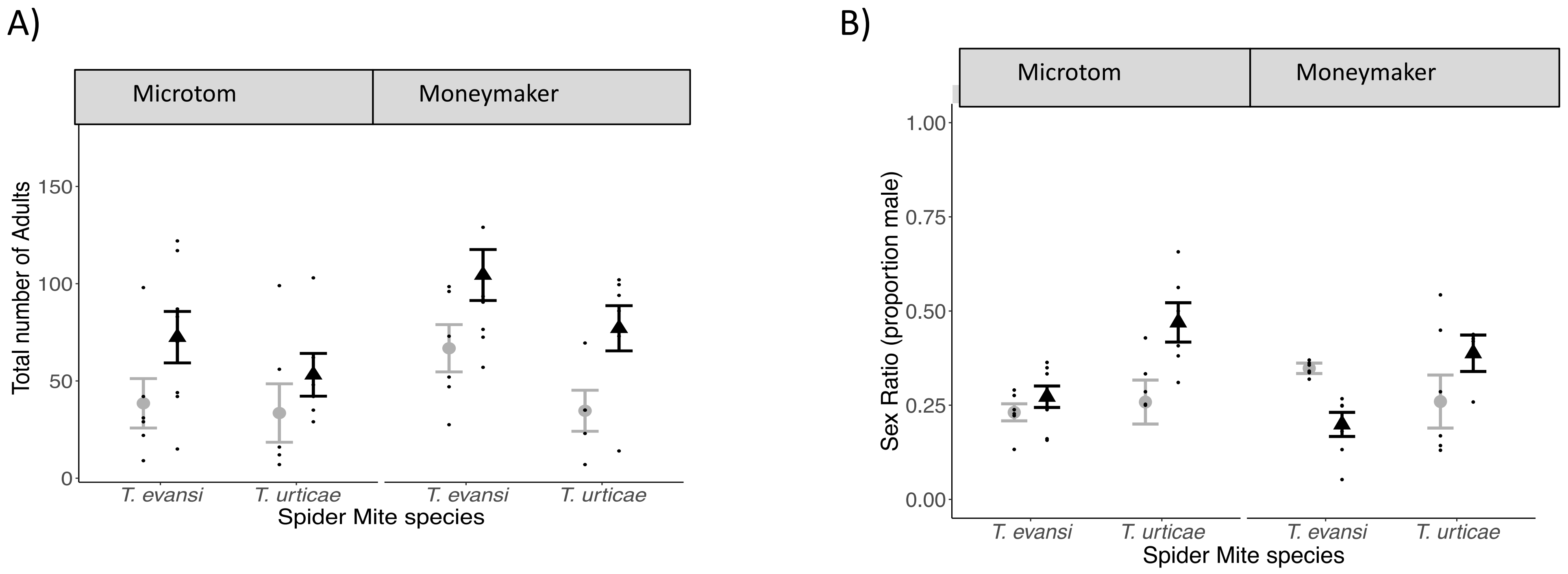


Figure S5: Effect of Tomato spotted wilt virus on mean (± standard error) *Tetranychus urticae* and *Tetranychus evansi* A) adult offspring, B) offspring sex ratio on mock (gray) and virus infected (black) plants of the Microtom and Moneymaker varieties. Mean is represented by a gray dot in all non-TSWV treatments, and by a black triangle in all TSWV treatments. All the data points are jittered in black along the respective treatments.

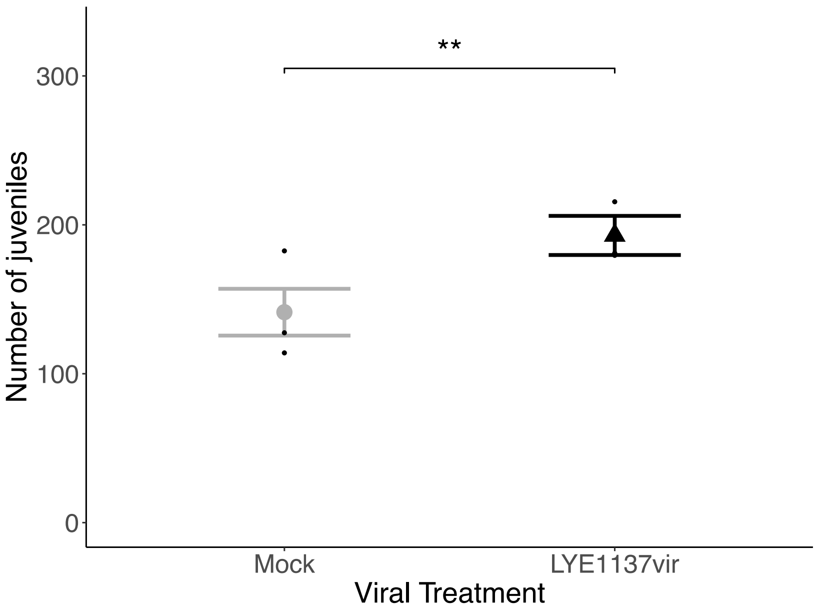


Figure S6: Effect of Tomato spotted wilt virus on mean (± standard error) *T. urticae* A) number of juveniles on Moneymaker whole plants mock infected (gray) or infected with TSWV strains France81 and LYE1137vir (black). ** represent the significant differences between both the viral treatments. Mean is represented by a gray dot in all non-TSWV treatments, and by a black triangle in all TSWV treatments. All the data points are jittered in black along the respective treatments.

**A1: Distribution of TSWV in infected tomato plants.**

**Methods:** In a separate experiment, we investigated TSWV spread in infected plants 11 days after infection. For this, we infected 15 Moneymaker plants, 5 each with the virus strains France 81, LYE 1137vir, LYE 55 as described in the main text. We then checked all leaves of infected plants using Agdia test strips on day 11 instead of just taking a sample from the topmost leaves. We used a GLMM with a binomial error structure in glmmTMB to investigate if there was an effect of leaf position, viral strain and their interaction on the probability of being infected. The leaf that was inoculated with the virus was denoted as zero and the leaf below as -1 and immediately above +1, 2, 3 and 4. Virus strain was included in the model as fixed a factor and leaf as a covariate. Individual plant replicate was nested within virus strain in the model as a random factor.

**Results:** There was a significant effect of viral strain ($\chi^2_2$ = 14.19, p = < 0.001) and leaf number ($\chi^2_5$ = 18.476, p = < 0.001) on the probability for leaves to be positive for TSWV infection, but no interaction between them ($\chi^2_2$ = 0.1075, p = 0.947). The two topmost leaves on the plant (leaves 3 and 4) always tested positive for infection, whereas leaves further down the plant were mostly negative. Plants infected with viral strains LYE 1137vir and France 81 were more likely to have leaves positively infected lower down the plant (including the leaf that was inoculated) than viral strain LYE 55 (Figure S6).

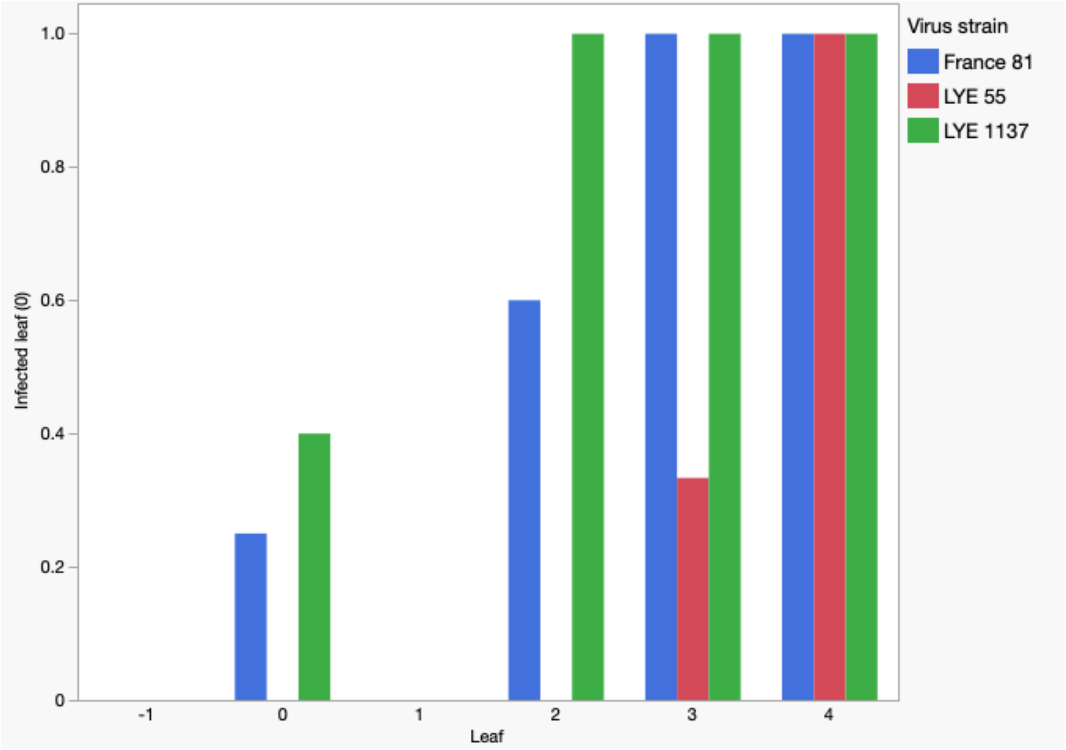


Random effects in the model:
Virus strain(plant): 1.617e-09 ± 4.021
Virus strain: 1.025e-11 ± 3.202e-06

Figure S7: Proportion of plants with leaves at different levels of the plant positive for infection with TSWV.
